# Heterozygous mutation of *Vegfr3* decreases renal lymphatics but is dispensable for renal function

**DOI:** 10.1101/2021.01.17.427041

**Authors:** Hao Liu, Chitkale Hiremath, Quinten Patterson, Saumya Vora, Zhiguo Shang, Andrew R. Jamieson, Reto Fiolka, Kevin M. Dean, Michael T. Dellinger, Denise K. Marciano

## Abstract

**Background:** Lymphatic abnormalities are observed in several types of kidney disease, but the relationship between the renal lymphatic system and renal function is unclear. The discovery of lymphatic-specific proteins, advances in microscopy, and available genetic mouse models provide the tools to help elucidate the role of renal lymphatics in physiology and disease.

**Methods:** We utilized a mouse model containing a missense mutation in *Vegfr3* (dubbed *Chy*) that abrogates its kinase ability. *Vegfr3*^*Chy*/+^ mice were examined for developmental abnormalities and kidney-specific outcomes. Control and *Vegfr3*^*Chy*/+^ mice were subjected to cisplatin-mediated injury. We characterized renal lymphatics using a combination of tissue clearing, light-sheet microscopy and computational analyses.

**Results:** In the kidney, we found Vegfr3 is expressed not only in lymphatic vessels, but also various blood vessels. *Vegfr3^Chy/+^* mice had severely reduced renal lymphatics with 100% penetrance, but we found no abnormalities in blood pressure, renal function and histology. Similarly, there was no difference in the degree of renal injury after cisplatin, although *Vegfr3^Chy/+^* mice developed more perivascular inflammation by histology. Control mice treated with cisplatin had a measurable increase in cortical lymphatic density despite no change in cortical lymphatic volume and length.

**Conclusions:** We demonstrate that Vegfr3 is required for development of renal lymphatics, but a reduction in lymphatic density does not alter renal function and induces only modest histological changes after injury. Our data suggests that an increase in lymphatic density after cisplatin injury may reflect the loss of cortical volume associated with chronic kidney disease rather than growth of lymphatic vessels.

**SIGNIFICANCE STATEMENT:** Defects in renal lymphatics occur in various kidney diseases, but their role in maintaining kidney structure and function is unknown. We combine tissue clearing, light-sheet microscopy and computational analysis to characterize lymphatics and find that mice with a heterozygous mutation in *Vegfr3* (*Vegfr3*^*Chy*/+^) have severely reduced renal lymphatics. Strikingly, these mice have indistinguishable renal function and histology compared with controls. Even after cisplatin injury, there are no differences in renal function, although *Vegfr3^Chy/+^* mice developed more perivascular inflammation. Our data present a novel method of lymphatic quantification and suggest that a normal complement of renal lymphatics is dispensable for renal structure and function.

## INTRODUCTION

The recent discovery of proteins uniquely enriched in lymphatic endothelia, combined with the increasing capabilities of mouse genetics, have ushered in new insights on the function of the lymphatic system. Lymphatic vessels serve as conduits for the transport of extracellular fluid, immune cells, and macromolecules, but like their blood counterparts, lymphatic endothelial cells (LECs) are remarkably heterogenous, demonstrating organ-specific functions in homeostasis and disease (1–3).

Several types of kidney disease have been associated with renal lymphatic abnormalities including polycystic kidney disease, hypertensive nephropathy and cisplatin-induced kidney disease, among others (4–6). Acute disruption of renal lymphatics in animal studies by surgical ligation demonstrates variable effects on renal function and structure (7, 8). Additionally, human primary lymphatic disorders have been associated with renal abnormalities, but these defects are not well characterized (9). Thus, despite their associations with disease, the role of lymphatics in kidney development, physiology and disease remains unclear.

The growth of lymphatics or lymphangiogenesis is mediated in part by vascular endothelial growth factors c and d (Vegf-c/d) acting on the receptor Vegfr3 (10, 11) to promote LEC proliferation, migration, and survival (2, 3, 12). Indeed, administering Vegf-c or Vegf-d has been shown to increase renal lymphatic density in experimental mouse models (6, 13). Prior studies have shown that mice null for *Flt4*, the gene encoding Vegfr3 (hereafter denoted *Vegfr3*), have embryonic lethality due to cardiovascular defects, while heterozygous mice appear phenotypically normal (14). Primary human lymphedema (Milroy’s disease) is a rare autosomal dominant disorder caused by a missense mutation in the tyrosine kinase domain of *Vegfr3* that results in hypoplastic dermal lymphatics (15); however, it is not known if visceral and specifically, renal lymphatics are affected by this mutation.

Renal lymphatics are generally confined to the renal cortex, where they are found adjacent to medium and large sized blood vessels (16, 17). The low relative density of lymphatics and the lack of lymphatic-specific markers in the kidney makes quantification and assessment of lymphatic changes challenging. Some well-characterized lymphatic markers, such as Lyve-1, Podoplanin, Vegfr3, and Prox1, are present in non-lymphatic cells within the kidney (17–19). To address these limitations, a recent study generated three-dimensional (3D) reconstructions of lymphatic vessels in developing mouse and first-trimester human kidneys (20). Use of 3D methods in adult mouse models will improve our ability to identify and quantify lymphatics, and thus increase our knowledge of how lymphangiogenesis affects renal outcomes in experimentally-induced models of renal disease.

In the current study, we utilized a mouse model carrying a mutation in *Vegfr3* (dubbed *Chy*) that renders it kinase inactive (21) and examined its effect on the development of renal lymphatics. We developed a computational workflow to quantitatively assess renal lymphatics in 3D using cleared, immunostained kidneys that were imaged with isotropic spatial resolution using axially swept light-sheet microcopy (22). Our results shed new light on the role of renal lymphatics in physiology and disease, and present a rigorous and more precise method for analyzing and quantifying lymphatic changes.

## MATERIALS AND METHODS

### Animals

We bred *Vegfr3*^*Chy*/+^ mice (C3H101H-Flt4Chy/H, from MRC Harwell) (21) to generate mutant and heterozygous mice and crossed *Vegfr3*^*Chy*/+^ mice to *Prox1-tdTomato*^Tg/+^ mice (23) to generate wildtype and Chy mice with the Prox-1 reporter. Mice were maintained on mixed genetic backgrounds consisting largely of C3H and B6;129S, and there was 100% penetrance of the reduced renal lymphatic phenotype in *Vegfr3*^*Chy*/+^ mice. Mice were genotyped by standard PCR. Neonatal kidneys were harvested and fixed for 2 hours in 4% paraformaldehyde (PFA) in phosphate-buffered saline (PBS). Adult mice were perfused with PBS and then 4% PFA, and kidneys fixed for 4 hours.

### Cisplatin Injury

Age matched (8-9 week old) male and female mice were administered low dose (5mg/kg), medium dose (10mg/kg), high dose (15mg/kg) cisplatin by two intraperitoneal injections two weeks apart adapted from Landau et al. (24). Cisplatin (Sigma Aldrich, 15663-27-1) was dissolved to 1mg/ml in sterile 0.9% normal saline. Mice were sacrificed and kidneys were harvested four weeks after initial injection. Procedures were performed according to UTSW IACUC-approved guidelines.

### Biochemical Measurements

Serum Cr was measured by HPLC (UTSW O’Brien kidney research core).

### Blood Pressure Measurements

Systolic and diastolic blood pressure was determined by tail cuff measurements using the Kent Scientific 53170 non-invasive CODA high throughput system. Mice were placed in pre-warmed chambers until tail temperatures reached > 34°C, and then 20-25 measurements were performed. Measurements were obtained at the same time each day for 5 days, but only data from the fourth and fifth days were used for analysis. Individual measurements were accepted if tail volume > 40μl.

### Immunofluorescence and Histology

Fixed kidneys were permeabilized with 0.3% Triton X-100/PBS (PBST) and blocked with 10% donkey sera/PBST as performed previously (25). Antigen retrieval was performed with Trilogy (Cell Marque). Samples were incubated with primary antibodies overnight (4°C), then with fluorophore-conjugated secondary antibodies. Kidney sections were mounted with Prolong Gold (Invitrogen). The following antibodies were used at 1:100 unless stated otherwise: Lyve-1 (Abcam, ab14917), Prox1 (Millipore, ABN278, RFP (Rockland, 6004010379), Vegfr3 (R&D systems, AF743), Endomucin (SCB, sc-65495), UTB (1:200, a gift of Jeff Sands, Emory U.), MECA-32 (Plvap) (SCB, sc-19603). Confocal imaging was performed on a Zeiss LSM880 confocal microscope. Images were minimally processed and re-sampled to 300 dpi using Adobe Photoshop. Histologic stains, including Periodic acid-Schiff and Masson’s trichrome, were performed by the UTSW O’Brien kidney research core.

### Quantification of Perivascular Inflammation

Mid-kidney transverse sections were stained with Periodic acid-Schiff and the number of medium sized artery and vein pairs in the cortex (interlobular arteries and veins) were manually counted per section. Perivascular inflammation was defined as the presence of greater than or equal to 2 layers of mononuclear inflammatory cells surrounding vessels.

### Light-sheet Imaging and Quantification

Fixed kidneys were sectioned using a vibratome then prepared using the CUBIC method (26). Sections were immersed in CUBIC-L at 37 °C. After delipidization, samples were washed and incubated with primary antibodies followed by fluorophore-conjugated secondary antibodies. Samples were incubated in CUBIC-R+ for refractive index matching prior to imaging with cleared tissue axially swept light-sheet microscopy (27). Quantitative image analysis was performed using MATLAB (*Mathworks, R2019b*). 3D reconstructions were generated by computationally fusing image sub-volumes with BigStitcher (28). IMOD image analysis software was used for 3D rotation and structure cropping (29). Projected images were rotated to measure the tissue boundary along the z-axis to estimate tissue thickness. To detect the edge of the tissue, 3D projections were enhanced using a built-in MATLAB function ‘histeq’ and a threshold (>mean density) was used to measure the edge of the tissue. Similarly, the edge of the lymphatic tree was detected by setting a separate threshold (0.4*mean density) to generate a binary map. The lymphatic edge shape then was used to automate a medullary mask and an arcuate mask to separate the cortical lymphatics and arcuate lymphatics. We then isolated the region between the tissue edge and the arcuate mask. The area of this region is then multiplied by the tissue thickness to calculate the cortical volume. The medullary and arcuate masks were combined, and the image projection was then extended to 3D to demarcate segmentation boundaries of the lymphatic structures. The arcuate mask was manually tuned to select the optimal distance from the tissue edge to separate the cortical lymphatics from the arcuate lymphatics. A built-in MATLAB function (bwareaopen.m) was used to remove noise from the background. The volume of lymphatic structures was calculated by summing the number of voxels of lymphatics in the 3D binary map. To calculate total length of the lymphatic vasculature, skeletonization was performed using a medial surface/axis thinning algorithm (30). The skeleton maps included both a surface skeleton and an axis skeleton so we had to develop separate algorithms to calculate each prior to summation. A convolution function was used to detect the location of the surface and axis skeletons. The length of the 3D surface skeleton (Ls) was calculated using equation (1). Di is the diameter of each surface skeleton, which was generated by doubling the maximum distance of all points in same class to their center position.

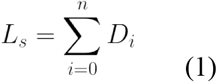

To calculate the length of the 3D axis skeleton (L_a_), equation (2) was adapted from a previously reported method using ImageJ (31). Nsingle is the number of grids set to 1 with all neighboring pixels (i.e., with distance < 2) set to 0. D_ij_ is the distance (< 2) between two neighboring grids.

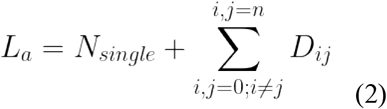

### Statistics

All data shown are mean ± SD. Statistical significance was performed using unpaired, two-tailed Student’s t-test or an unpaired, nonparametric (Mann-Whitney) test as indicated. Survival curves were evaluated with the log rank test. All statistical analysis was performed with GraphPad Prism 8 (San Diego, CA).

## RESULTS

### Vegfr3 localizes to renal lymphatics and blood capillaries in adult mouse kidneys

Previous studies have shown that Vegfr3 localizes to blood and lymphatic endothelia during development but becomes restricted to lymphatic endothelia in adulthood except in some specialized capillary beds (32). Immunolocalization of Vegfr3 in neonatal (P0) mouse kidneys demonstrated a high intensity Vegfr3 signal colocalizing with the lymphatic marker Lyve-1 (17) (Fig. 1A, arrow). Low intensity Vegfr3 was present in blood endothelial cells (Fig. 1A, arrowhead). In adult mouse kidney, high levels of Vegfr3 were present in cortical renal lymphatics. Surprisingly, discernable levels of Vegfr3 were also present in blood endothelia of peritubular capillaries, although the blood endothelia of medium-sized arteries and veins had little-to-no Vegfr3 (Fig. 1B). High magnification images demonstrated that Vegfr3 is present in blood capillary networks extending from the renal cortex to the medulla, including the ascending and descending vasa recta (Fig. 1C). These results demonstrate that Vegfr3 expression is not specific to lymphatic endothelia in the kidney.

**Figure 1.**
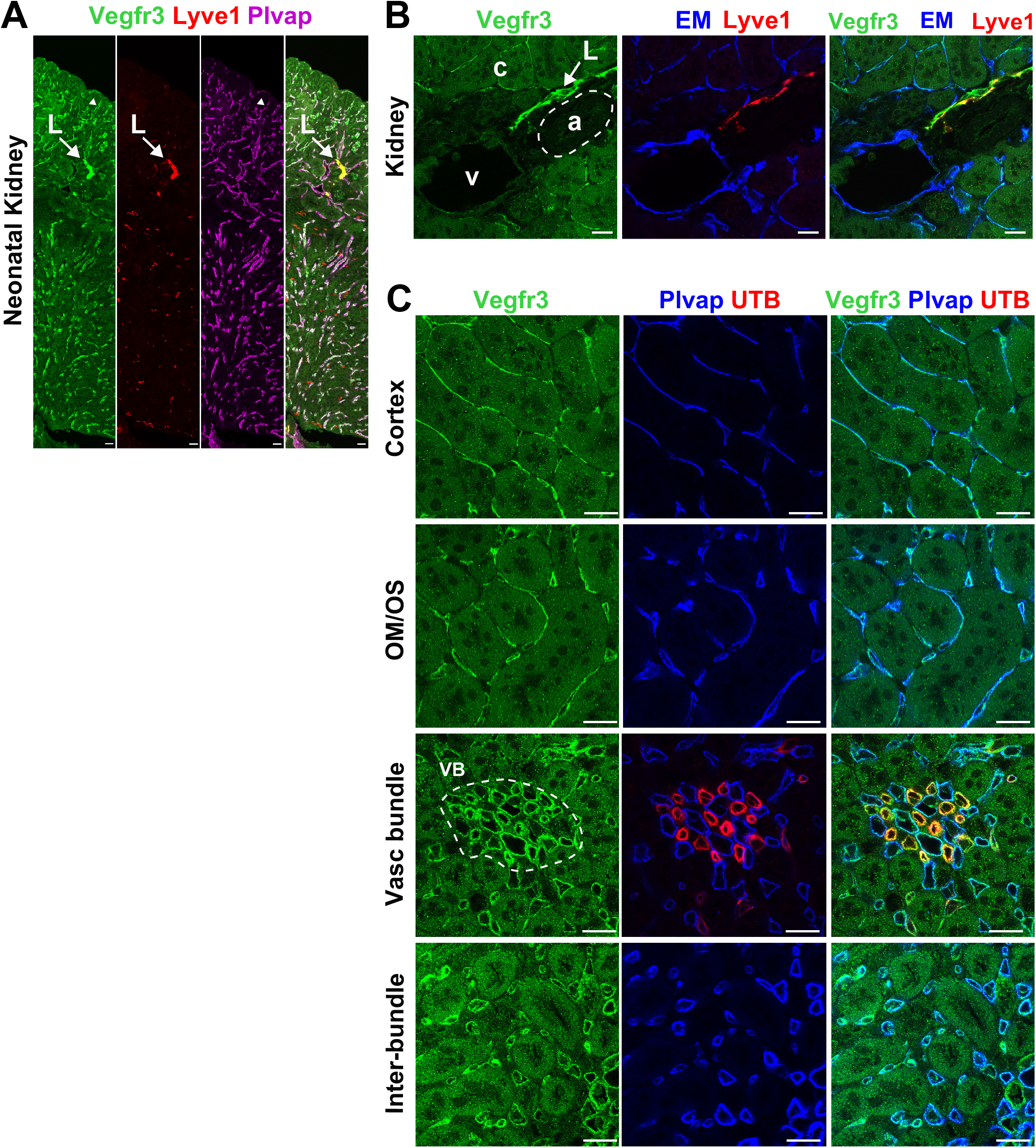
Vegfr3 localizes to lymphatic and select blood endothelia in neonatal and adult mouse kidneys. (A) Immunolocalization of Vegfr3 in neonatal mouse kidney labels lymphatics (L, arrow) marked by Lyve-1 and also co-localizes with blood endothelia (arrowhead) marked by Plvap (purple). (B) Vegfr3 expression in adult mouse kidney is highest in lymphatics (L) but also present at lower levels in surrounding capillaries (c). There is absence of Vegfr3 in medium sized arteries (a, white dotted line) and veins (v). (C) Immunolocalization of Vegfr3 in various anatomical capillary beds throughout the adult mouse kidney including the peritubular capillaries and the ascending and descending vasa recta in the vascular bundle (VB) marked by Plvap (blue) and UTB (red), respectively. Scale bars: 20 μm in (A, B and C). OM/OS, Outer Stripe of Outer Medulla. Results are representative of four independent experiments.

### Loss of Vegfr3 kinase function results in absent renal lymphatics

We examined embryonic kidneys (e17.5) immunostained with Lyve-1 from *Vegfr3*^+/+^, *Vegfr3*^*Chy*/+^, and *Vegfr3^Chy/Chy^* mice. The kidneys were cleared and then imaged using axially swept lightsheet microscopy (see methods). The 3D image data of homozygous mutant (*Vegfr3^Chy/Chy^*) kidneys revealed a complete absence of renal lymphatic vasculature (Fig. 2). Heterozygous mice had reduced renal lymphatics that primarily localized to the renal hilum, suggesting a delay or deficit in lymphatic development. Importantly, the *Vegfr3*^*Chy*/+^ mice did not appear to have a global delay in development (not shown). In contrast, *Vegfr3^Chy/Chy^* mice were smaller than littermate controls and died in utero or during the neonatal period, precluding assessment of later time points. The non-lymphatic staining of Lyve-1 in embryonic kidneys is due to Lyve-1 expression in F4/80^+^ embryonic macrophages. Lyve-1 is absent from mature macrophages, as previously reported (17).

**Figure 2.**
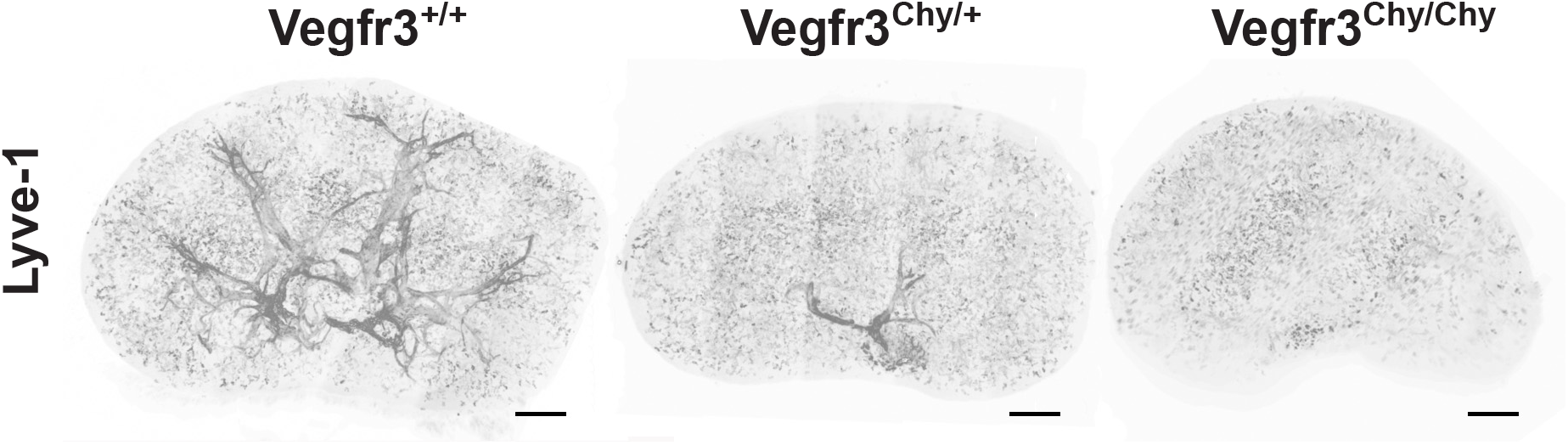
Missense mutation in Vegfr3 leads to reduced renal lymphatics during embryonic development. 3D reconstruction of renal lymphatics marked by Lyve-1 in e17.5 whole mouse kidneys of *Vegfr3^+/+^, Vegfr3^Chy/+^*, and *Vegfr3^Chy/Chy^* mice. Non-lymphatic, punctate labeling of Lyve-1 represents individual macrophages. Scale Bar: 100 μm. Results are representative of two independent experiments.

### Heterozygosity of *Vegfr3* mutation reduces renal lymphatic volume, length and density

We first developed a workflow to qualitatively and quantitatively assess renal lymphatics (Fig. 3). We bred *Vegfr3*^*Chy*/+^ mice to mice carrying a transgenic Prox1-tdTomato allele. Prior studies have demonstrated that mice carrying this allele have tdTomato expression in lymphatics (23). Kidneys were harvested from the mice at 12 weeks of age, sectioned transversely (1 mm), and cleared. Kidney sections were then imaged using axially swept light-sheet microscopy (22, 27). Approximately 3000-4000 images (each 870.4μm x 870.4μm x 999.6μm) were stitched together (28) to create a 3D reconstruction of each tissue sample.

**Figure 3.**
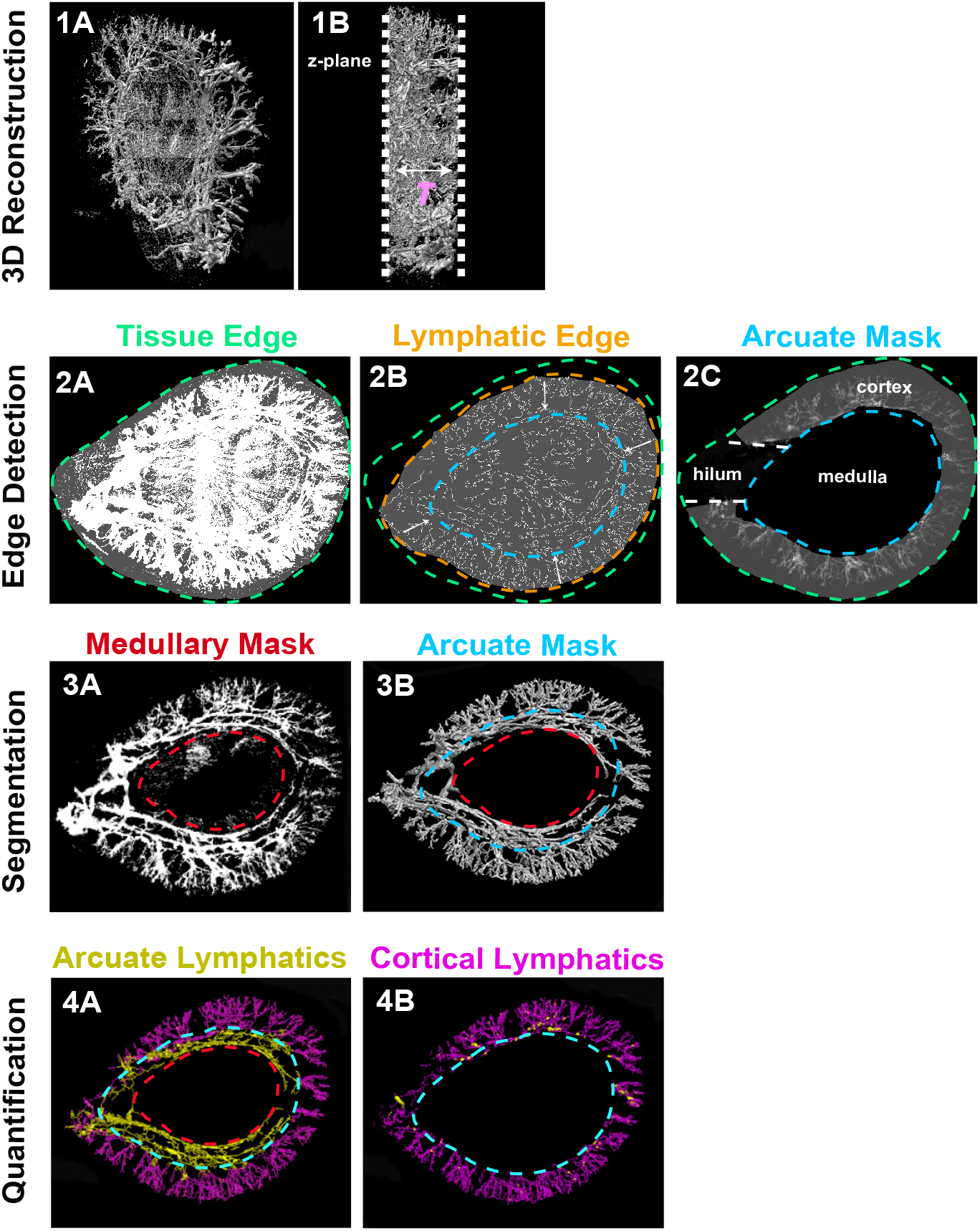
Workflow of renal lymphatic quantification and analysis. (1A) Composite image from raw image stacks after integrating a series of 3D sub-tiles. (1B) is the view after rotating the image perpendicular to the z-axis to estimate tissue thickness (*T*). (2A and 2B) Separate intensity thresholds are used to detect the edge of the tissue (green) and the edge of the cortical lymphatic structures (orange). The shape of the lymphatic structure edge is then used to generate an arcuate mask (blue) and a medullary mask (red). (2C) Cortical volume is calculated by isolating the region between the arcuate mask and the tissue edge. (3A and 3B) 2D image projection with masks is extended to 3D. (4A and 4B) Medullary and arcuate masks are manually tuned to segment the arcuate (yellow) and cortical (purple) lymphatics. Quantification of volume is done by summing the number of voxels in lymphatics in the 3D binary map. A 3D skeleton map is generated using a thinning algorithm to calculate lymphatic length (see methods).

After generating a composite 3D image of the whole tissue (Fig. 3, 1A and 1B), a 2D projection along the z-axis was performed on the entire volume to generate an image used for determining a mask that segments the lymphatic structures from the rest of the tissue. Using separate intensity thresholds, we outlined the edges of the entire tissue (Fig. 3, 2A) and the periphery of the lymphatic tree (Fig. 3, 2B). The shape of the lymphatic structure edge was used to generate an ‘arcuate mask’ and when combined with the tissue edge, allowed us to calculate the renal cortical volume (Fig. 3, 2C). A second mask was applied to remove the non-lymphatic Prox1 signal in the medulla (Fig. 3, 3A). The 2D masks were then extended to 3D (Fig. 3, 3B) and superimposed to separate the cortical (yellow) and arcuate (purple) lymphatics (presented as binary maps in Fig. 3, 4A and 4B). Based on these cortical and arcuate binary maps, we then calculated total lymphatic volume, total lymphatic length, cortical lymphatic volume, and cortical lymphatic length (see methods). We normalized lymphatic volume and length to both total kidney volume and cortical volume to obtain total lymphatic density and cortical lymphatic density, respectively.

**Figure 4.**
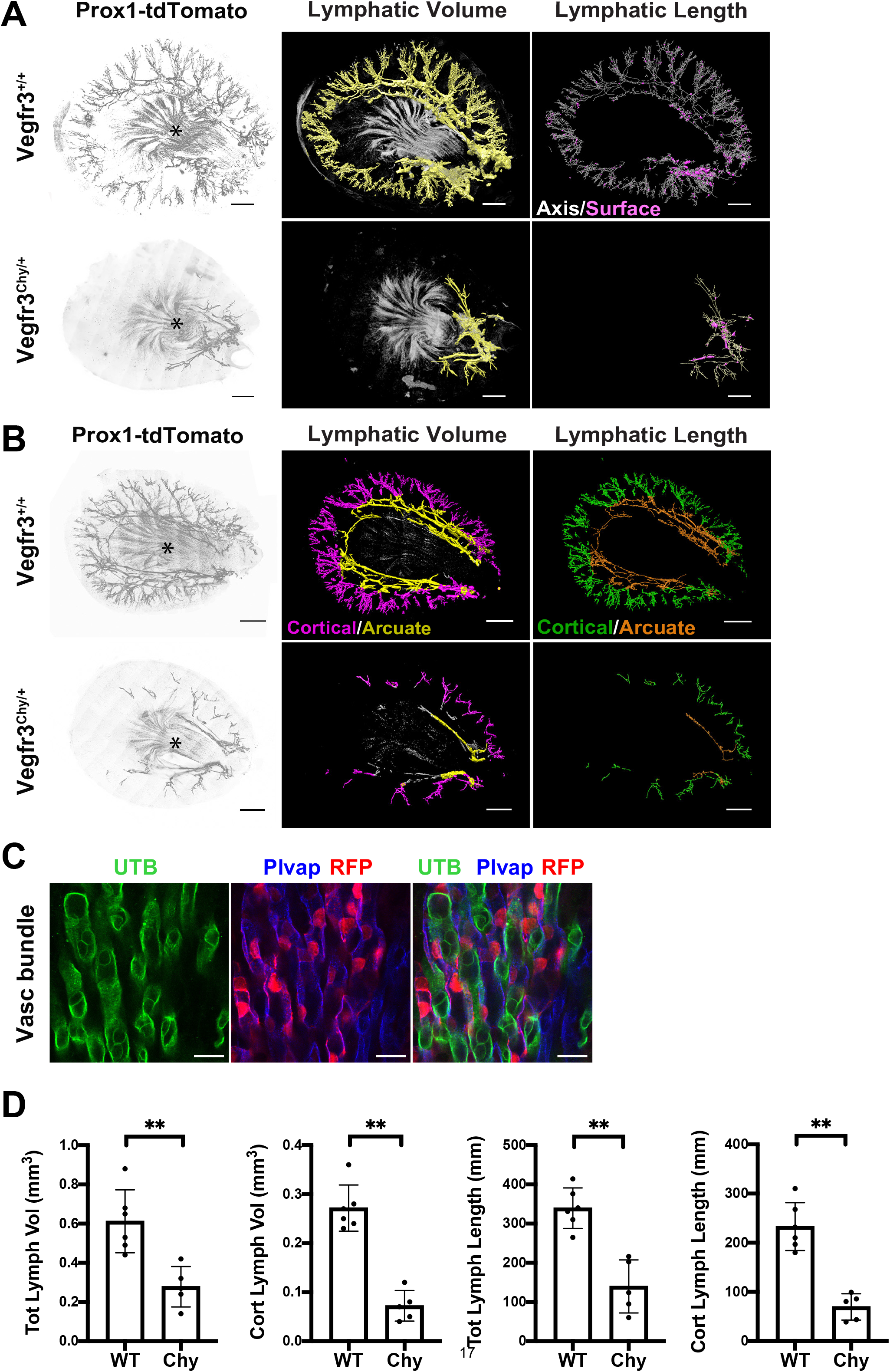
*Vegfr3^Chy/+^* mice have decreased renal lymphatic volume and lymphatic length. (A) 3D image data of 12-week-old *Vegfr3^+/+^* (control) and *Vegfr3*^*Chy*/+^ (Chy) kidneys carrying a *Prox1-tdTomato* reporter allele. Left panel - grayscale rendering, Middle panel - total lymphatic volume projection, Right panel - skeletonized lymphatic length projection. Axis skeleton and surface skeleton are summed to calculate the total length of lymphatics. See methods for details. (B) 3D image data of a separate pair of control and *Vegfr3*^*Chy*/+^ kidneys with volume and length projections segmented by cortical lymphatics (purple and green) and arcuate lymphatics (yellow and orange). Asterisks in (A and B) show non-lymphatic *tdTomato* signal in medullary vascular bundles. (C). Immunolocalization of *tdTomato-* expressing cells in ascending vasa recta (Plvap, blue) but not descending vasa recta (UTB, red). Results are representative of three independent experiments. (D) Quantification of total lymphatic volume, cortical lymphatic volume, total lymphatic length, and cortical lymphatic length in control and *Vegfr3^Chy/+^* kidneys. See methods for details. (\*\**P<0.005*). Analyzed by Mann-Whitney test. Scale bars: 1 mm in (A and B); 20 μm in (C).

Our initial experiments characterized the localization of tdTomato in *Prox1-tdTomato*^*Tg*/+^ mice. We discovered that the *Prox1-tdTomato^Tg/+^* allele delineated a lymphatic tree in *Vegfr3*^+/+^ (control) mice that predominantly resides in the cortex and appears to follow the major blood vessels of the kidney (Fig. 4A and 4B, row 1), consistent with prior anatomical descriptions of renal lymphatics (33). We applied the same image processing workflow to evaluate adult *Vegfr3*^*Chy*/+^ mice and characterized the severe reduction in renal lymphatics. This finding demonstrated that the defect in *Vegfr3*^*Chy*/+^ embryonic mice (Fig. 2) persisted into adulthood (Fig. 4A and 4B, row 2). The tdTomato signal was also present in the renal medulla where renal lymphatics are absent (Fig. 4 A-B, asterisks). In the medulla, a specialized blood endothelial capillary network, the ascending and descending vasa recta, is responsible for interstitial fluid recycling and uptake (34). Through immunolocalization studies, we determined that medullary tdTomato co-localized with the ascending vasa recta (AVR), but not descending vasa recta (DVR) (Fig. 4C). This extra-lymphatic expression in the AVR has been previously described in another Prox1 reporter mouse model (18).

To quantify the reduction in renal lymphatics observed in *Vegfr3*^*Chy*/+^ mice, we measured lymphatic volume and length as described previously. We found that absolute values of total lymphatic volume, total lymphatic length, cortical lymphatic volume and cortical lymphatic length were reduced in *Vegfr3*^*Chy*/+^ mice compared with *Vegfr3*^+/+^ littermate controls (Fig. 4D). The decrease in renal lymphatics was 100% penetrant in the *Vegfr3*^*Chy*/+^ mice. When corrected for kidney size, total lymphatic volume/total kidney volume and total lymphatic length/total kidney volume were still reduced (Fig. S1A). Of note, the total kidney volume (the denominator) was not different between groups (Fig. S1B). Cortical lymphatic density, as approximated by cortical lymphatic volume/total cortical volume and cortical lymphatic length/total cortical volume, appeared to be even more dramatically reduced in *Vegfr3*^*Chy*/+^ mice (Fig. S1C).

### Reduced lymphatics do not affect systemic blood pressure or cause renal insufficiency

Recent studies have suggested a link between renal lymphatics and systemic blood pressure (4, 35–37). To determine the effects of attenuated renal lymphatics on blood pressure, we measured tail cuff pressures on control and *Vegfr3*^*Chy*/+^ mice at 8-9 weeks of age. There was no significant difference in systolic blood pressure or diastolic blood pressure between the two groups (Fig. 5A). To evaluate the effects of reduced lymphatic volume and length on renal function, we performed biochemical measurements to measure serum creatinine. We found no differences in serum creatinine (Fig. 5B). Furthermore, histological evaluation did not reveal overt differences in glomerular morphology, tubular morphology or degree of fibrosis. Renal insufficiency did not develop over time as mice aged to 9 months showed no differences in serum creatinine (Fig. 5B) or histology (Fig. S2A and S2B).

**Figure 5.**
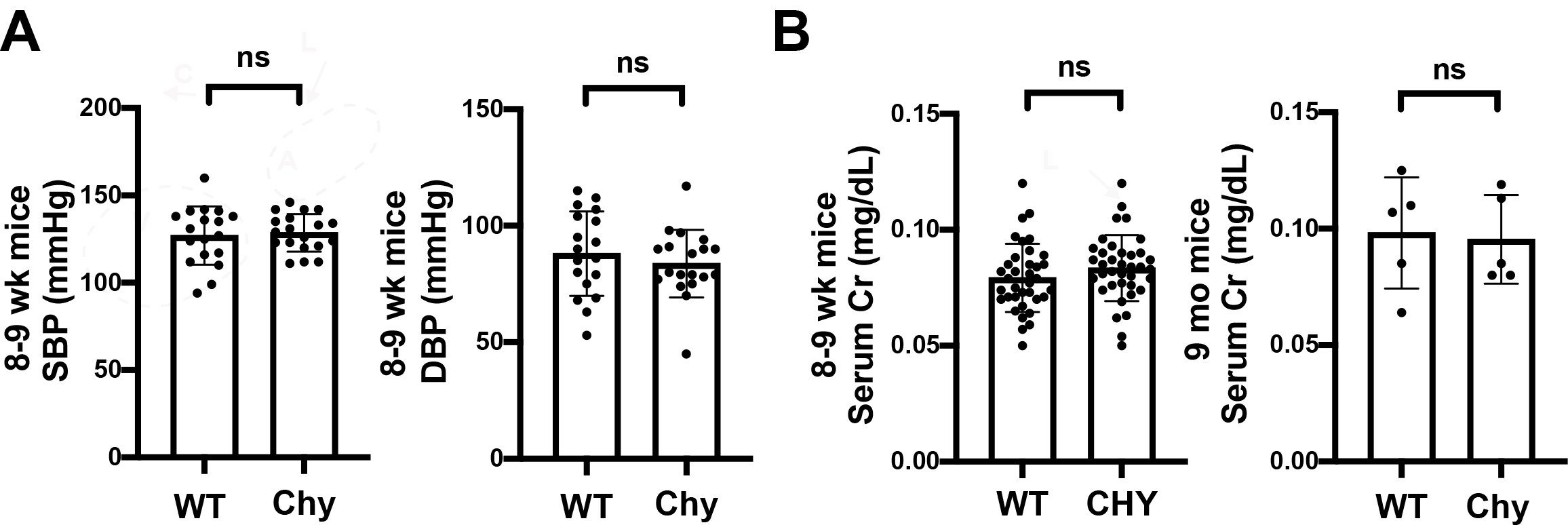
*Vegfr3^Chy/+^* mice do not develop blood pressure abnormalities or kidney disease over time. (A) Tail cuff measurements of systemic blood pressures from 8-9 week old wild type (WT, *Vegfr3^+/+^;* n = 18) and mutant (‘Chy’, *Vegfr3^Chy/+^*; n = 19) mice. (B) Measurements of renal function by serum creatinine from 8-9 week old mice (*Vegfr3*^+/+^; n = 36; *Vegfr3^Chy/+^*; n = 37) and 9 month old mice (*Vegfr3*^+/+^; n = 5; *Vegfr3^Chy/+^*; n = 5). Analysis with unpaired *t* test. SBP, systolic blood pressure; DBP, diastolic blood pressure.

### Reduced lymphatics do not affect kidney outcomes after cisplatin-induced chronic kidney injury

Next, we investigated the effect of reduced renal lymphatics on renal function following cisplatin-induced kidney injury. Murine models have shown that administering two cisplatin doses two weeks apart result in sustained loss of glomerular filtration rate (GFR), and thus cause chronic kidney disease (24). Because of the heterogeneity of cisplatin dosing protocols and its unknown effect on survival of *Vegfr3*^*Chy*/+^ mice, we designed experimental arms administering vehicle (0.9% saline), low (5mg/kg), medium (10mg/kg), or high dose (15mg/kg) cisplatin. The 8-9 week-old *Vegfr3^+/+^* and *Vegfr3*^*Chy*/+^ mice were administered two intraperitoneal doses of cisplatin or vehicle two weeks apart, and mice were sacrificed 4 weeks after the first injection (Fig. 6A). Serum creatinine was measured prior to the first injection and at the 4-week endpoint.

**Figure 6.**
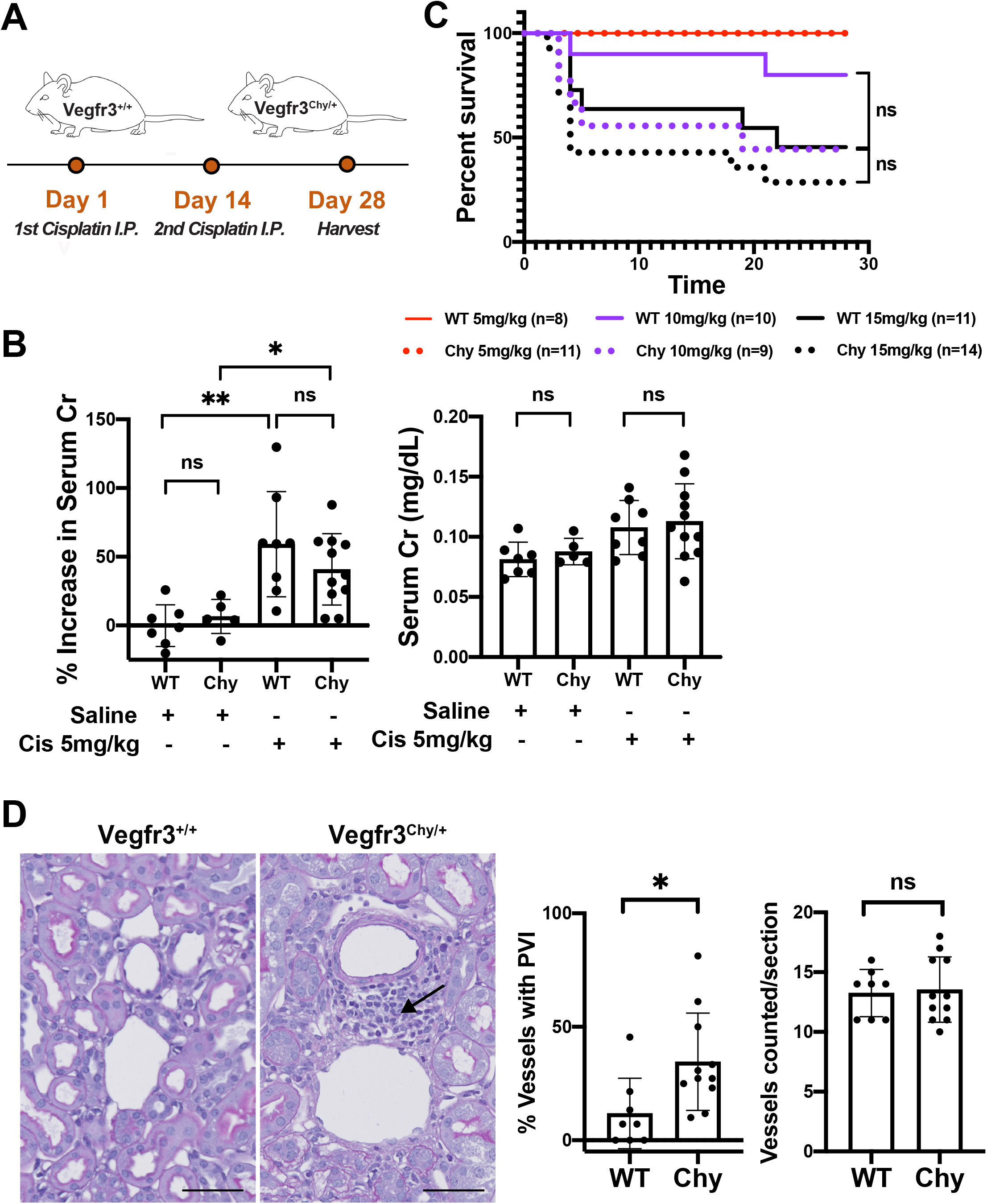
*Vegfr3^Chy/+^* mice are not more susceptible to cisplatin-induced chronic kidney disease. (A) Schematic of cisplatin dosing regimen given two doses, two weeks apart. (B) Percent increase in serum creatinine and comparison of renal function after low dose cisplatin vs saline injection. (C) Survival curves for low (5 mg/kg), medium (10 mg/kg), or high dose (15 mg/kg) cisplatin groups. (D) Histology and quantification of increased perivascular inflammation (arrow) in *Vegfr3*^*Chy*/+^ mice compared to control. See methods for details (\**P<0.05, **P<0.005*). Analyzed by Mann Whitney test. Scale bars: 50 μm in (D).

All mice that received cisplatin had increased serum creatinine and histological abnormalities suggestive of kidney disease (Fig. 6B and S3A). Some mice that received medium or high doses of cisplatin did not survive to the 4-week endpoint. Specifically, in the high dose group, 5/10 (50%) of wild-type and 10/14 (71%) of *Vegfr3^Chy+^* mice died before 4 weeks. In the medium dose group, 2/9 (22%) of wild-type vs and 4/9 (44%) of *Vegfr3*^*Chy*/+^ mice died. Kaplan-Meier curves for these experimental groups showed no statistical differences in survival between *Vegfr3^+/+^* and *Vegfr3*^*Chy*/+^ mice at 4 weeks (Fig. 6C). Measurements of serum creatinine (or percentage change in serum creatinine) at 4 weeks were not different between the genotypes in mice that received 10 or 15mg/kg cisplatin (Fig. S3A); however, this result should be interpreted with caution because of the high mortality rate in these mice.

All mice that received low dose (5mg//kg) cisplatin or vehicle survived. Mice in the low dose cisplatin group developed an increase in serum creatinine over 4 weeks compared with mice that received vehicle only. Surprisingly, there was no difference in the magnitude of renal dysfunction observed between wild-type and *Vegfr3*^*Chy*/+^ mice in the low dose cisplatin group, as assessed by serum creatinine (Fig. 6B). However, histologic evaluation revealed increased mononuclear inflammatory cells surrounding medium-sized vessels in *Vegfr3*^*Chy*/+^ mice, although this did not translate into worse renal outcomes (Fig. 6D).

### Cisplatin-induced CKD increases cortical density of lymphatics but not total cortical lymphatic volume or length

To evaluate the effect of cisplatin-induced kidney injury on renal lymphangiogenesis, we performed light-sheet microscopy on 1 mm transverse sections from cisplatin-treated (10mg/kg, 2 doses) and vehicle-treated *Prox1-tdTomato^Tg/+^* mice. The mice were age- and sex-matched (3 males and 3 females for each arm).

We generated 3D reconstructions of the sections using the endogenous tdTomato signal (Fig. 7A). As before, we masked the medullary signal and delineated cortical and arcuate lymphatic vessels. This was necessary because the arcuate vessels were often not fully visualized in a section and thus, had the potential to introduce bias (Fig. 7A, white dotted line). Additionally, it is cortical lymphangiogenesis that has been correlated with clinical outcomes in kidney disease (38). The cisplatin treated mice selected for imaging all developed renal dysfunction as evidenced by an increase in serum creatinine compared to the controls (Fig. 7B and S4A).

**Figure 7.**
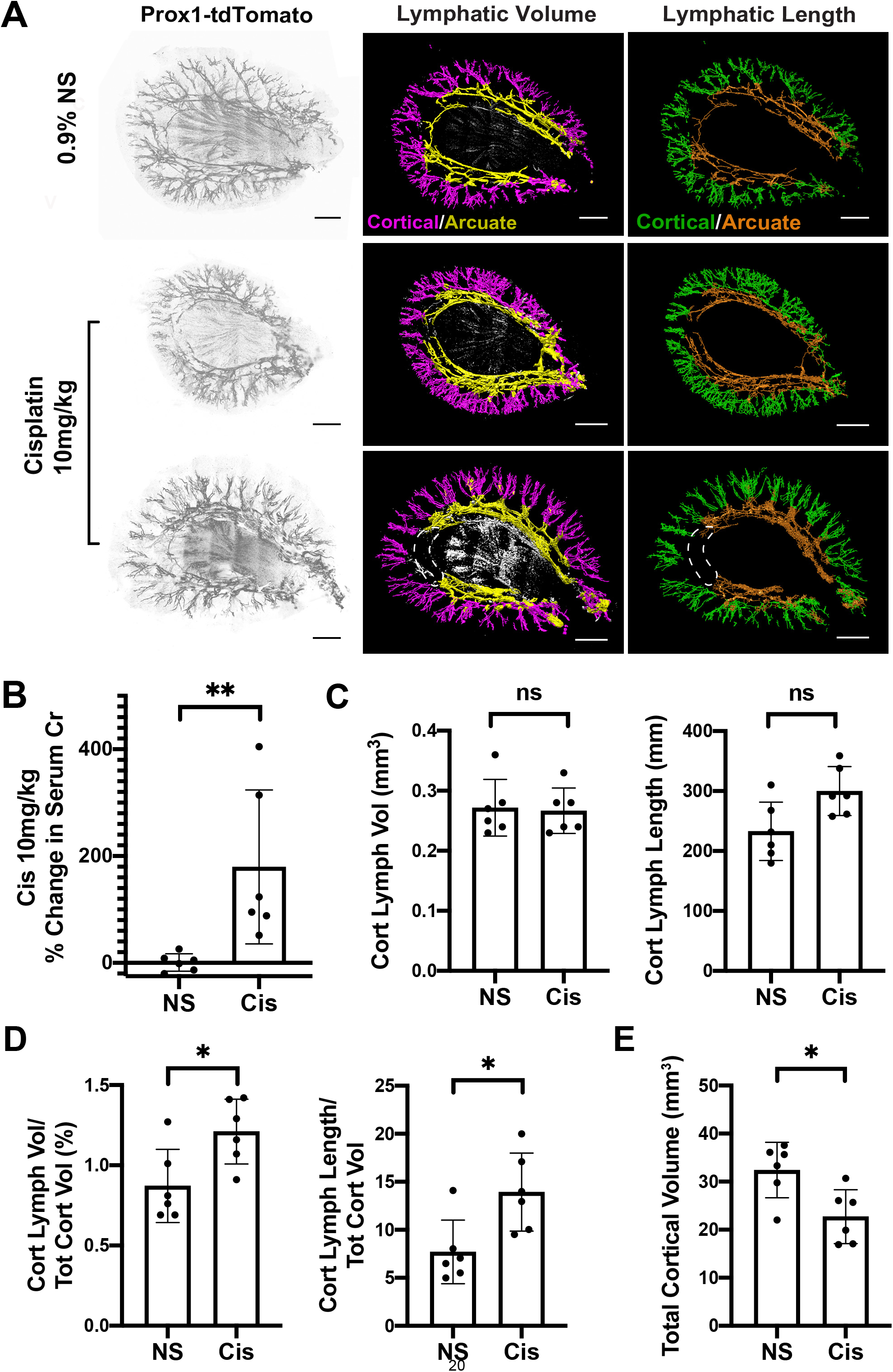
Cisplatin-induced CKD increases renal lymphatic density in the cortex due to loss of cortical volume. (A) 3D reconstructions of *Vegfr3^+/+^* (control) kidneys carrying a *Prox1-tdTomato* reporter allele after saline injection or 10 mg/kg cisplatin. Left panel – grayscale rendering, Middle panel – lymphatic volume projection, Right panel – skeletonization and lymphatic length projection. Renal lymphatics separated by anatomical location (cortical – purple/green; arcuate – yellow/orange). White dotted line outlines partially missing arcuate lymphatics not visualized in this sample. (B) The kidneys selected for imaging in the 10mg/kg cisplatin group developed significant kidney injury as measured by a % increase in serum creatinine. (C) Comparison of absolute values of cortical lymphatic volume and cortical lymphatic length between groups. (D) Quantification cortical lymphatic density measured by cortical lymphatic volume (purple)/total cortical volume and cortical lymphatic length (green)/total cortical volume. (E) Cisplatin treated mice have a reduction in cortical volume compared to vehicle. See methods for details (\**P<0.05, **P<0.005*). Analyzed by Mann Whitney U test. Scale bars: 1 mm in (A).

We found that the total volume and length of cortical lymphatics was not different in the vehicle and cisplatin-treated groups (Fig. 7C). However, the cortical density of lymphatics (as approximated by the cortical lymphatic volume/total cortical volume and cortical lymphatic length/total cortical volume) was increased in cisplatin-treated mice compared with controls (Fig. 7D). Importantly, the numerators were not significantly different, but total cortical volume was decreased in cisplatin-treated mice (Fig. 7E and S4B), a finding that occurs commonly in many types of chronic kidney disease. Baseline mouse weights prior to injections were not different between the two groups (Fig. S4C).

## DISCUSSION

The role of renal lymphatics in maintaining kidney homeostasis and the lymphatic response to kidney injury is unclear. In this study we used light-sheet microscopy combined with tissue clearing to demonstrate that a heterozygous mutation in *Vegfr3* leads to significantly decreased renal lymphatics. Importantly, the reduced lymphatics do not affect systemic blood pressure, renal function, or worsen cisplatin-mediated chronic kidney disease. Additionally, we applied novel 3D imaging and post-processing techniques to measure lymphatic changes after cisplatin injury and found that lymphatic density increases in the renal cortex as a result of reduced cortical volume, not due to a change in lymphatic volume or length.

Our immunolocalization studies demonstrate that Vegfr3 is not a specific marker for renal lymphatics, a finding corroborated by single cell RNA-seq datasets (39). The functional significance of Vegfr3 expression in renal blood capillaries is not clear. Although we did not perform an in-depth assessment, we did not observe gross blood vascular abnormalities, and systemic blood pressure was unchanged in *Vegfr3*^*Chy*/+^ mice. Vegf-c/d are ligands that bind to Vegfr3 to regulate lymphangiogenesis, and pharmacologic treatment or genetic manipulation of either has protective effects on renal function in various types of kidney disease (6, 37). Our finding of Vegfr3 expression in multiple capillary beds throughout the kidney suggest that therapeutic intervention of Vegf-c or Vegf-d may affect blood endothelia, possibly promoting angiogenesis and/or survival in a larger capacity than previously expected. Thus, the protective effects of these pro-lymphangiogenic factors may not be solely due to lymphatic changes in the kidney.

The 3D reconstruction of embryonic kidneys with a homozygous *Vegfr3* mutation revealed a complete absence of renal lymphatics, while a heterozygous mutation resulted in attenuated lymphatic growth and reduced lymphatic density that persisted into adulthood. We expected that a reduction in renal lymphatics would cause an inability to adequately drain interstitial fluid, thereby leading to parenchymal swelling and secondarily reducing renal function over time. Surprisingly, we found that *Vegfr3*^*Chy*/+^ mice had normal renal clearance, estimated by serum creatinine, and no histological evidence of interstitial edema or fibrosis. The idea that disrupting renal lymphatic drainage does not affect long-term renal survival is not new. The ligation of donor kidney lymphatics prior to renal transplantation shows that human kidneys can function long term after surgery (40). Several studies have shown that disrupting renal lymphatics in animal models leads to enlarged kidneys but found no interdependence between renal lymphatic drainage and kidney function (8, 41, 42). However, not all studies had similar findings (7).

One possible explanation for why we did not see interstitial edema in *Vegfr3*^*Chy*/+^ mice is that the low lymphatic density was sufficient to meet drainage demands in homeostasis. Another possibility is the presence of intrinsic, compensatory mechanisms. Apart from lymphatics, interstitial fluid can exit the kidney via venous and urinary routes. Studies have shown that renal lymph is derived from both glomerular capillary filtrate and reabsorbed tubular fluid (43, 44). One can postulate that changes in hydrostatic and oncotic pressure gradients caused by reduced lymphatic density could lead to increased venous and/or urinary efflux of fluid. This hypothesis is supported by early studies in animal models that show ligation of renal lymphatics induces a diuresis (8, 42, 45). Future studies will be needed to examine possible renal hemodynamic and tubular compensatory mechanisms in the *Vegfr3* mouse model.

Renal lymphangiogenesis has been reported to occur in many different clinical scenarios, including cisplatin-induced kidney disease, amongst others (4, 5, 38, 46, 47), and it has generated a logical enthusiasm for targeting the lymphatic system in a wide spectrum of kidney diseases (6, 36, 37, 48). Cisplatin injury did not lead to differences in renal outcomes in *Vegfr3*^*Chy*/+^ mice. However, it should be noted that our analysis focused on the low dose cisplatin group because of the mortality that occurred with higher doses. An average of 40% increase in serum creatinine in our low dose group may have not been severe enough to detect a difference in outcomes. Additionally, fibrosis and inflammation were not prominent histological findings after four weeks after injury (24). Thus, future studies that examine later time points with this injury model may be warranted.

Departing from conventional methods that manually count lymphatics per tissue section, we used a novel 3D quantification strategy to measure lymphatic changes after cisplatin injury. We found that cortical lymphatic density (cortical lymphatic volume and length / total cortical volume) increased after 4 weeks; however, the absolute values of total cortical lymphatic volume and length were not significantly different between mice treated with vehicle vs cisplatin. Rather, the calculated increase in cortical lymphatic density is the function of a decrease in total cortical volume, a frequent characteristic of chronic kidney disease. It is important to note that our results do not argue for or against the presence of renal lymphangiogenesis after cisplatin treatment. It is possible that lymphatic growth and regression many coexist and we speculate that remodeling does occur in injury states. Our results demonstrate that evaluating changes in tissue volume is critical for obtaining accurate changes in lymphatic (and potentially blood) angiogenesis in kidney disease models.

There is still much to learn about the role renal lymphatics play in kidney health and whether or not they exacerbate or defend against certain types of renal injury. What is clear, however, is that contemporary methods of counting lymphatics in slide sections to quantify changes in lymphatic density may not accurately portray what is occurring at the organ level. This is especially true in the context of chronic kidney disease where reduced cortical volume is an inevitable outcome.

## AUTHOR CONTRIBUTIONS

Conceptualization: DKM, HL, MTD; Supervision: DKM; Data analysis: HL and DKM; Writing of the first draft: HL, DKM. All authors performed experiments, generated figures and reviewed/revised the manuscript.

## ACKNOWLEDGEMENTS

The C3H101H-Flt4Chy/H (Vegfr3) mice were obtained from the MRC Harwell which distributes this strain on behalf of the European Mouse Mutant Archive (EMMA: www.infrafrontier.eu). The repository number is EM:00068.

## SUPPLEMENTAL TABLE OF CONTENTS

1. **Figure S1.** *Vegfr3*^*Chy*/+^ mice have decreased total renal lymphatic density and decreased cortical lymphatic density.

2. **Figure S2.** *Vegfr3*^*Chy*/+^ mice have no gross histological abnormalities after 9 months.

3. **Figure S3**. No differences in the degree of kidney injury after medium and high doses of cisplatin.

4. **Figure S4.** Cisplatin-mediated kidney injury induces renal injury and decreases kidney volume.

---

**Figure S1.**
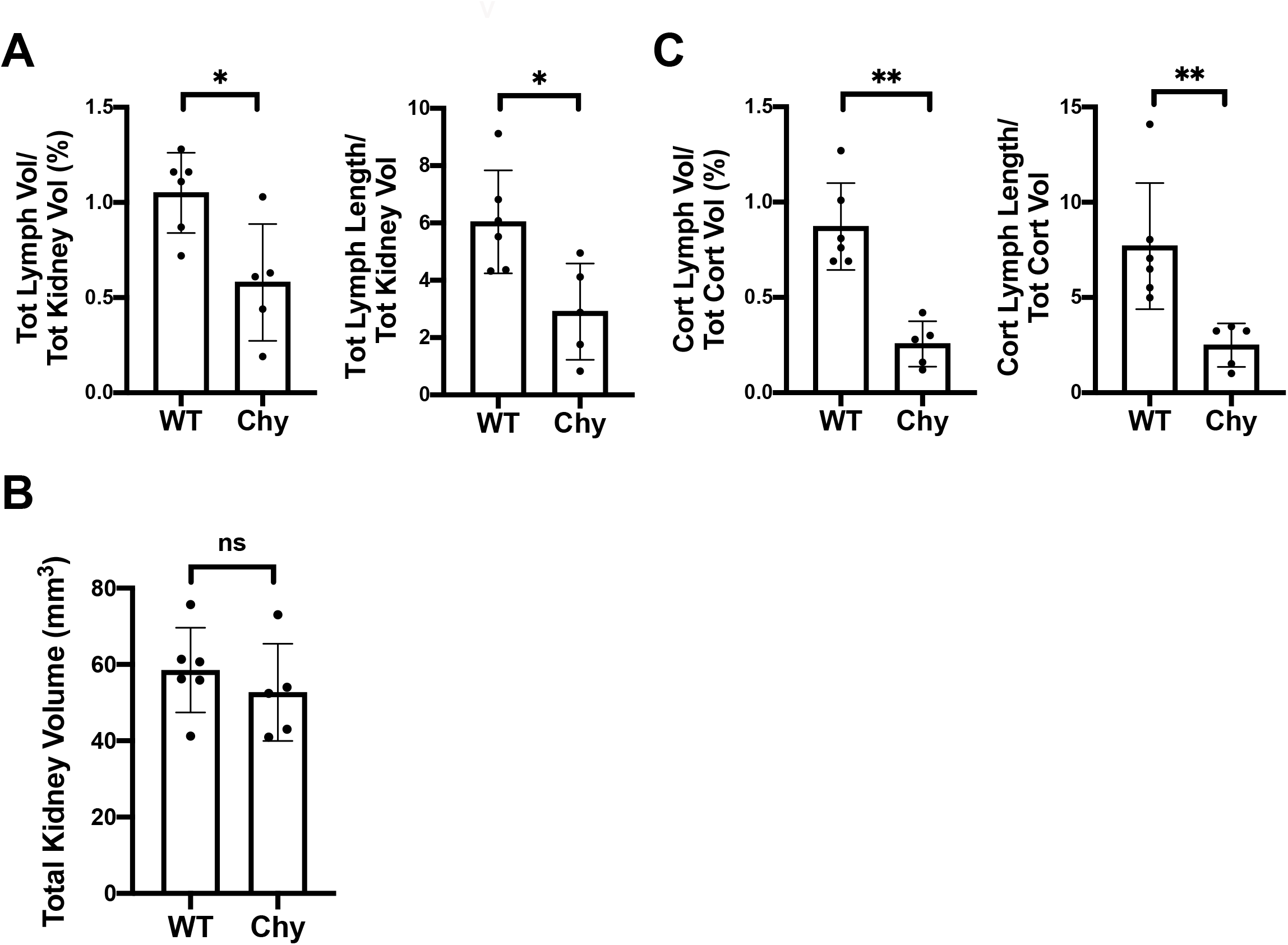
*Vegfr3*^*Chy*/+^ mice have decreased total renal lymphatic density and decreased cortical lymphatic density. (A) Quantification of renal lymphatic density measured by total lymphatic volume/- total kidney volume and total lymphatic length/total kidney volume. (B) Quantification of total kidney volume in *Vegfr3^+/+^* (control) and *Vegfr3*^*Chy*/+^(*Chy*) kidneys. (C) Quantification of cortical lymphatic density measured by cortical lymphatic volume/total cortical volume and cortical lymphatic length/total cortical volume.

**Figure S2.**
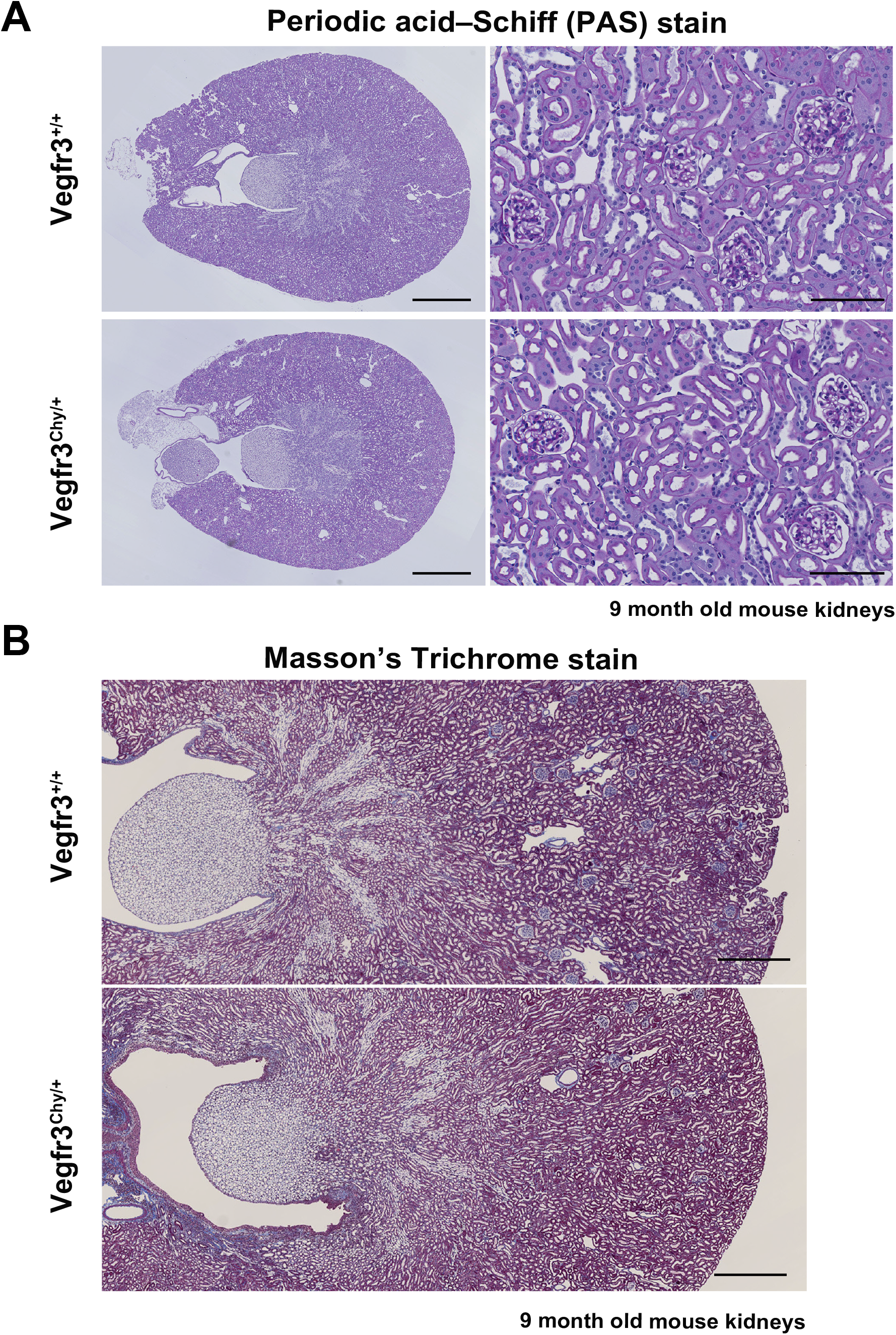
*Vegfr3*^*Chy*/+^ mice have no gross histological abnormalities after 9 months. (A) Periodic acid Schiff stain in *Vegfr3*^+/+^ (control) and *Vegfr3*^*Chy*/+^(*Chy*) mice shows no significant glomerular or tubular defects. (B) Masson’s Trichrome stain shows no significant difference in renal fibrosis. Results are representative images obtained from five mice of each genotype.

**Figure S3.**
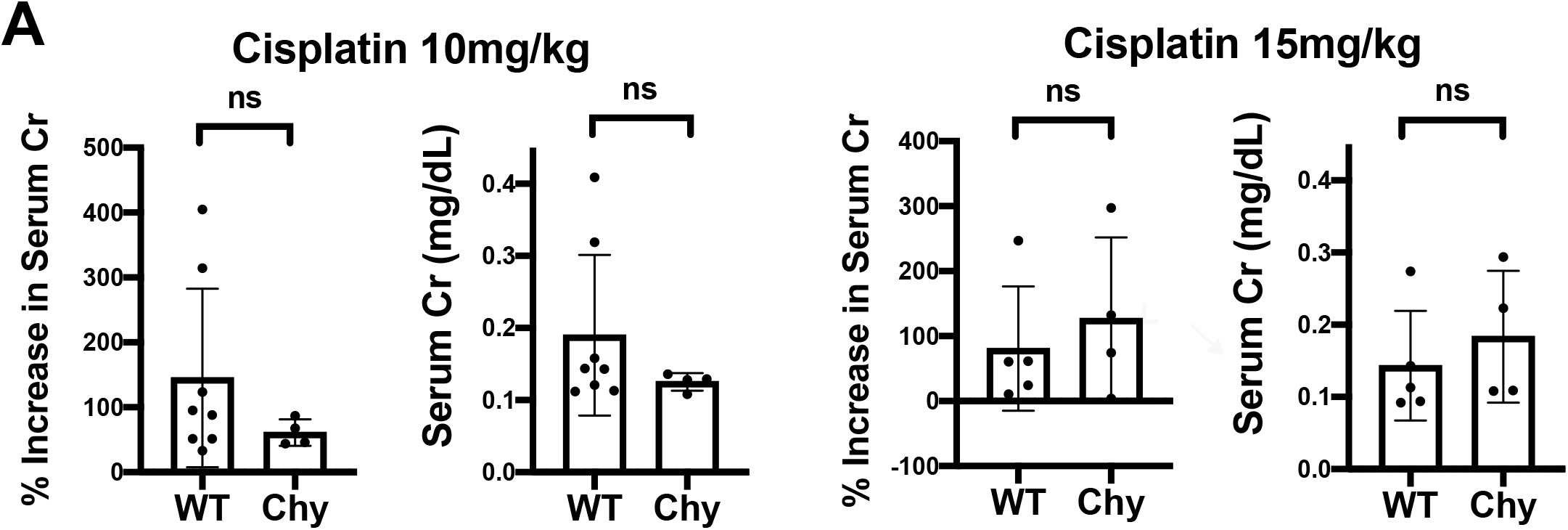
No differences in the degree of kidney injury after medium and high doses of cisplatin. (A) Percent increase in creatinine and comparison of renal dysfunction after medium (10mg/kg) and high (15mg/kg) dose cisplatin. This data is difficult to interpret given high early mortality rates and selection for survivors.

**Figure S4.**
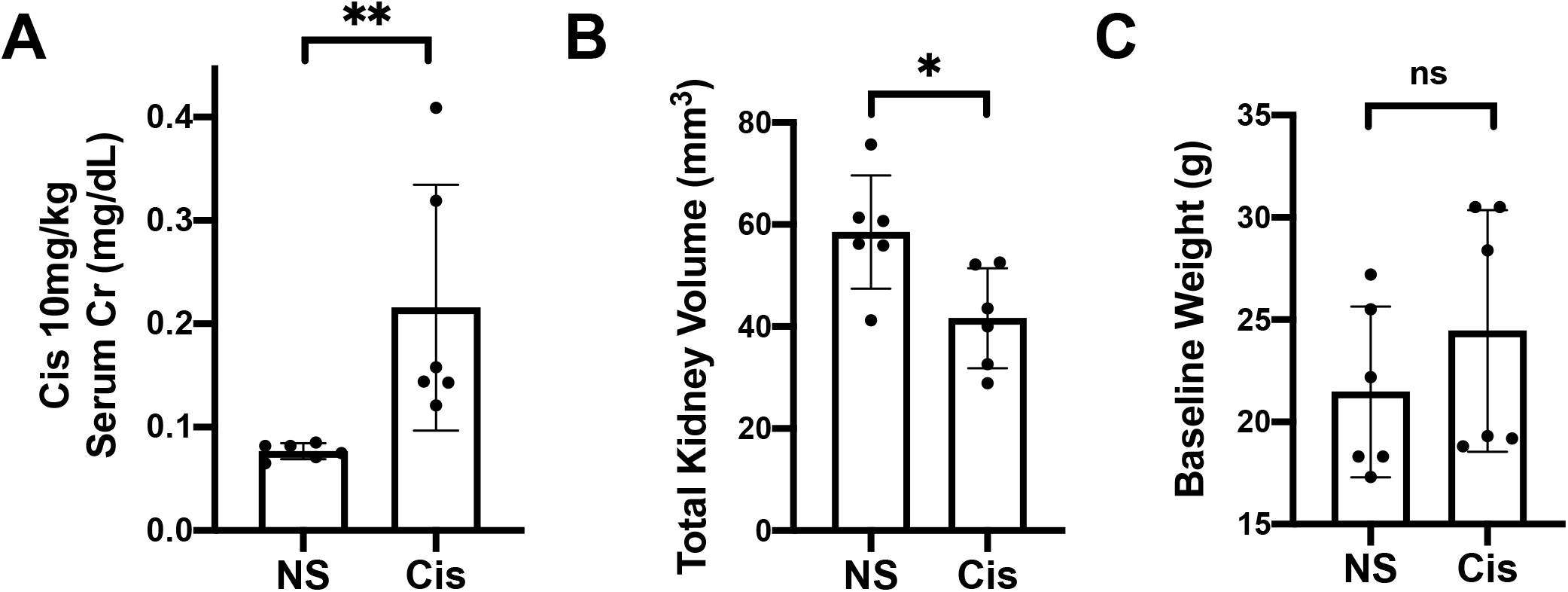
Cisplatin-mediated kidney injury induces renal injury and decreases kidney volume. (A) Measurements of serum creatinine at 4 weeks after two doses of cisplatin 10mg/kg. (B) Quantification of total kidney volume in the cisplatin-treated group compared to vehicle. (C) Animal weights in both groups before treatment.

